# Obesity and diabetes related disturbances in systemic metabolism induce a cardiometabolic heart failure with preserved ejection fraction phenotype in a combined high-fat, high sucrose diet and streptozotocin mouse model

**DOI:** 10.1101/2025.10.23.684124

**Authors:** Andre Heinen, Andre Spychala, Lucas Ballmann, Stefanie Gödecke, Zewa Faradj, Katharina Bottermann, Heba Zabri, Jens Fischer, Patrick Petzsch, Axel Gödecke

## Abstract

**Background:** Diabetes and obesity are associated with an increased incidence of heart failure with preserved ejection fraction (HFpEF), but the underlying pathophysiological mechanisms are poorly understood. A shortage of appropriate preclinical mouse models reflecting different pathophysiological disease aspects might contribute to this inadequate understanding of the complex and diverse HFpEF pathophysiology. We conducted a comprehensive analysis of a non-genetic, inducible T2DM mouse model with regard to its suitability as a preclinical model of cardiometabolic, diabetes-induced HFpEF.

**Methods:** T2DM was induced in C57Bl/6 mice by high-fat/high-sucrose diet (HFHSD) combined with low-dose streptozotocin (STZ) treatment (DIO-STZ), control animals received standard chow (chow). As additional control groups, animals of a DIO group were solely fed HFHSD throughout the study, and a STZ group received solely STZ injections while maintained on a standard chow. Cardiac function was assessed *in vivo* by echocardiography and left ventricular catheterization, as well as *in vitro* using the isolated perfused heart model. Structural, molecular and bioenergetic disturbances were analyzed by immunohistochemistry, RNA-seq, qPCR, western blot, and extracellular flux analysis of myocardial tissue, among others.

**Results:** Blood glucose, fatty acids and ketone body levels were elevated, and insulin plasma level were reduced in DIO-STZ animals compared to chow. DIO-STZ mice showed a strong cardiometabolic HFpEF phenotype with reduced cardiac output, end-diastolic volume, and increased filling pressure. Neither STZ nor DIO mice showed signs of HFpEF development. No difference in myocardial fibrosis nor in *in vitro* myocardial stiffness was detected between DIO-STZ and chow. Myocardial RNA-Seq clearly pointed towards disturbances in lipid and ketone metabolism. Extracellular flux analysis in intact cardiac tissue slices revealed an increased fatty acid oxidation capacity without differences in glucose metabolism. Mitochondrial respirometry revealed no indication of general mitochondrial dysfunction or mitochondrial uncoupling, but a reduced capacity for β-hydroxybutyrate oxidation.

**Conclusions:** The diabetic DIO-STZ mouse model showed a pronounced functional HFpEF phenotype. However, we show clear evidence that the underlying mechanism differs remarkably from the other HFpEF models making the DIO-STZ model a relevant extension of the range of HFpEF mouse models, especially for investigating molecular mechanisms or therapeutical interventions in diabetes associated HFpEF.

## Background

Diabetes mellitus is a group of metabolic disorders characterized by hyperglycemia, typically resulting from an absolute or relative insulin deficiency, which may arise from an unhealthy lifestyle, obesity, or genetic factors. The International Diabetes Federation (IDF) estimates that 537 million people worldwide are affected by diabetes, and this number is expected to rise to approximately 783 million by 2045 (1). Although high blood glucose levels in individuals with diabetes can now be relatively well managed through exogenous insulin administration, the comorbidities and long-term complications frequently associated with this complex disease continue to pose substantial challenges for affected individuals despite treatment (2).

Heart failure is one of the comorbidities increasingly diagnosed in individuals with diabetes. Up to 22% of people with diabetes show some form of heart failure, which can develop independently of risk factors such as coronary artery disease, hypertension, or valvular disease (3–6). The clinical incidence and mortality rates associated with heart failure in type 2 diabetes mellitus (T2DM) are significantly elevated −4 to 8 times higher than in non-diabetic patients - representing a growing burden on healthcare systems and economies, in addition to significantly diminishing the quality of life for affected individuals (6). This makes heart failure a significant cardiovascular complication in diabetes.

Nearly half of all heart failure cases are marked by left ventricular dysfunction, accompanied by preserved ejection fraction (HFpEF), with the incidence of HFpEF continuing to increase in contrast to heart failure with reduced ejection (HFrEF) (7, 8). Epidemiological data show that disturbances in systemic body metabolism as obesity, metabolic syndrome and/or T2DM are found in most patients with HFpEF suggesting these disturbances as major risk factors and pathophysiological drivers of a cardiometabolic HFpEF phenotype (9, 10). In addition, a causal significance of systemic metabolic deteriorations as obesity and diabetes in the development of HFpEF is supported by clinical trials showing that antidiabetic sodium-glucose cotransporter 2 inhibitors (SGLT2i) as well as obesity-reducing glucagon-like peptide-1 receptor agonists (GLP-1RA) have favorable effects in HFpEF patients (11, 12). Thus, HFpEF appears to be a systemic disease rather than a purely cardiac condition. Despite these robust clinical evidence that systemic metabolism can impact on HFpEF development, the understanding of the underlying mechanism resulting in the cardiac dysfunction is strongly limited, partially due to a lack of pre-clinical HFpEF animal models that mirror HFpEF features. It should be emphasized that, due to the heterogeneous pathophysiological facets of HFpEF development, it is unlikely that a single mouse model can comprehensively reflect all these facets of the disease. Therefore, developing specific animal models reflecting different pathophysiological HFpEF aspects is essential for a deeper understanding of the impaired mechanisms underlying each disorder, and for evaluating the efficacy of new therapeutics.

In this study, we investigate a diabetes mouse model based on a diet-induced obesity (DIO) due to high-fat high-sucrose diet (HFHSD) in combination with low dose streptozotocin (STZ) treatment modeling the transition from a pre-diabetic state to overt T2DM (13, 14). This model should reflect the clinically relevant T2DM disease development from adult-onset obesity to glucose intolerance, insulin resistance, the resulting compensatory insulin release and finally a partial pancreatic β-cell dysfunction/death. We demonstrate that the DIO-STZ mouse model develops a pronounced cardiac HFpEF phenotype with increased left ventricular filling pressures accompanied by robust alterations in myocardial substrate oxidation capacities towards increased fatty acid oxidation rates and reduced capacity to oxidize ketone bodies. These functional HFpEF changes are unique characteristics of the DIO-STZ model as they are neither detectable in mice that receive only STZ treatment nor in mice that solely were fed a HFHSD.

## Methods

### Animal procedures

C57BL/6 mice were purchased from Janvier Labs (Le-Genest-Saint-Isle, France), and were housed in groups of 2 per cage at constant room temperature on a 12-hour light/dark cycle with ad libitum access to water and food. The majority of the experiments were conducted using male mice of the C57Bl/6J substrain. In addition, to investigate potential sex or substrain dependent differences, female C57Bl/6J mice and male C57Bl/6N mice were used, respectively. All animals were allowed to acclimate after shipping for 1 week prior to the start of the experiment. All animal procedures were approved by the Animal Ethics Committee of the State Agency for Nature, Environment, and Consumer Protection North Rhine-Westphalia, Düsseldorf, Germany (file reference numbers 2024-267-Grundantrag, and 2024-276-Grundantrag), and were performed in accordance with the guidelines from Directive 2010/63/EU of the European Parliament. To induce diet-induced obesity (DIO), 9-week-old mice were fed a high-fat high-sucrose diet (HFHSD; 24% Sucrose, 60 kJ & Fat; ID S7200-E010; ssniff Spezialdiäten GmbH, Soest, Germany) for 11 weeks. After 5 weeks of feeding, animals received daily low-doses of streptozotocin (STZ; 50 mg/kg; i.p.) over five consecutive days to induce a partial pancreatic β-cell dysfunction. This group is referred to as DIO-STZ group; control animals received standard chow and no STZ treatment (chow). As additional control groups, animals of a DIO group were solely fed HFHSD throughout the study, and a STZ group received solely STZ injections while maintained on a standard chow.

### Plasma insulin, glucose and ketone body measurements as well as blood count

To measure blood glucose and beta-hydroxybutyrate levels, approximately 2 µl of blood was obtained by tail nicking, and measured using the StatStrip Xpress2 device. For blood cell count and measurement of plasma insulin levels, blood was collected via cardiac puncture and anticoagulated with 5 mmol/l EDTA. Blood cell count was performed using a VetScan® HM5 analyzer (ScilVet). To obtain blood plasma, collected blood was centrifuged at 2000×g for 20 minutes. The resulting plasma was analyzed for insulin concentration using the mouse insulin ELISA kit (DRG Instruments) following the manufacturer’s instructions.

### Quantitative determination of non-esterified fatty acids

The plasma concentration of non-esterified fatty acids (NEFA) was measured using the NEFA-HR(2) Assay (FUJIFILM Wako). A calibration line was constructed using NEFA Standard (0, 0.125, 0.25, 0.5 and 1 mM). Each plasma or standard sample (4 μl) was incubated with 200 μl of Fujifilm NEFA-HR(2) R1 for 5 minutes. The samples were then read using a microplate reader at a wavelength of 550 nm. Subsequently, 100 μl of Fujifilm NEFA-HR(2) R2 was added, incubated for an additional 5 minutes, and read again at 550 nm.

### Immunofluorescence staining

Pancreatic tissues samples were embedded in OCT embedding matrix (Cell Path Ltd, Newton, UK), snap-frozen at −40°C and cryo-sectioned at −22°C into 8 µm thick slices. Primary antibodies were incubated overnight at 4°C, followed by incubation with secondary antibodies for 3 hours at room temperature in the dark. The primary antibodies used were: Insulin (CS#8138, 1:100, mouse) and Glucagon (CS#2760, 1:200, rabbit). The secondary antibodies included Alexa488 goat anti rabbit (111-545-114) and Cy3 goat anti mouse (115-165-062), both obtained from Jackson ImmunoResearch Laboratories. The slides were mounted using DAPI Fluoromount-G (SouthernBiotech). Imaging and analysis were performed using a Keyence immunofluorescence microscope (BZ 9000) and ImageJ software.

### Echocardiography

Left ventricular function was evaluated using the Vevo2100 system (Visualsonics) with a 30 MHz linear transducer, following established protocols (15, 16). Images were captured at frame rates exceeding 200 frames per second. Mice were anesthetized with 2% isoflurane and positioned supine on a heated platform. ECG, respiration, and body temperature (maintained at 37±0.5°C with an infrared lamp as needed) were continuously monitored. The linear transducer was stabilized using a rail-mounted fixation system, and B-mode recordings of the parasternal long axis (PSLAX), as well as of the short-axis at mid-ventricular, apical, and basal level were collected. End-diastolic (EDV) and end-systolic volumes (ESV) were calculated by tracing the endocardial border in both diastole and systole. The Simpson method was applied for ventricular volume analysis, and ejection fraction (EF) was determined using the formula: EF = ((EDV - ESV) / EDV) * 100. Mid-ventricular SAX M-mode was used to determine diastolic thickness of the left ventricular posterior wall (LVPW) as well as the left ventricular internal diameter (LVID). Relative wall thickness (RWT) was calculated as twice the LVPW thickness divided by the LVID at end-diastole (LVPW × 2/LVID).

Speckle-tracking based strain imaging of PSLAX B-mode cine loops was used for more detailed analysis of ventricular function. Here, the global longitudinal strain (GLS), which represents a criterion within the HFA-PEFF score for HFpEF diagnosis (17), and the reverse peak of the global longitudinal strain rate (diastolic GLSR) as indicator of a diastolic dysfunction (18) were determined using the VevoStrain software package.

### Measurement of aortic and left-ventricular pressure

Aortic and ventricular pressures were determined in spontaneously breathing anesthetized mice (2% isoflurane, and 0.1 mg/kg buprenorphine) at a body temperature of 37±0.5°C. The catheter (PVR1035, Millar) was introduced into the aorta via the carotid artery to record the aortic pressure signal, and after advancing the catheter into the left ventricle (LV) the LV pressure signal was measured. Recordings were analyzed to obtain mean aortic pressure (AOP_mean_) as well as LV end-diastolic pressure (LVEDP) using Lab Chart Pro 8 software (ADInstruments).

### Analysis of cardiac fibrosis

After cervical dislocation, hearts were excised, fixed in 4 % paraformaldehyde, embedded in paraffin, and cut in 5 µm transverse heart sections. Total collagen of the Sirius Red (Polyscience, Inc, Warrington, PA) stained sections was measured under circularly polarized light. For this, five images (septum, anterior, lateral and posterior wall as well as right ventricular wall) from each section at 20x magnification were obtained using a Keyence BZ-X800 microscope. Colour images were converted to 8-bit images, and subsequently, collagen was quantified using a consistent threshold in a semi-automated manner using Image J software (version 1.54).

### Isolated perfused heart experiments

Mice received intraperitoneal injections of 250 IU heparin. After cervical dislocation, the heart was rapidly excised and transferred for preparation of the aortic trunk to ice-cold Krebs-Henseleit buffer. The aorta was cannulated, and the heart was perfused in non-recirculating Langendorff mode at constant pressure (80 mmHg) with a modified Krebs-Henseleit solution containing 118 mmol/L NaCl, 4.7 mmol/L KCl, 1.2 mmol/L MgSO_2_, 1.2 mmol/L KH_2_PO_4_, 25 mmol/L NaHCO_3_, 0.5 mmol/L EDTA, 2.25 mmol/L CaCl_2_, 8.32 mmol/L glucose, 1 mmol/L lactate, and 0.1 mmol/L pyruvate, equilibrated with 95% O_2_ and 5% CO_2_ (pH 7.4, 37°C) as described before (15). A fluid-filled balloon was inserted into the left ventricle, and ventricular preload was applied by setting end-diastolic pressure to 3-4 mmHg. After a stabilization period of 20 minutes, ventricular compliance was assesses by gradually increasing ventricular preload by stepwise increasing the volume of the balloon (2 µl per step; 20 steps), and the course of LVP_min_ was detected. In addition, dobutamine stress test was performed to investigate contractile reserve of DIO-STZ and chow hearts. Here, increasing concentrations of dobutamine (0, 10, 20, 50, and 100 nmol/l) were administered via the Krebs-Henseleit solution.

### RNA isolation and cDNA synthesis for RT-PCR

Tissues were homogenized using a Tissue Ruptor (Qiagen, Hilden) according to manufacturer’s protocol. Total RNA was extracted using Qiagen RNeasy columns, which included on-column DNase digestion. The quality of the isolated RNA was assessed using a microvolume spectrophotometer (Nanodrop™, Thermo Fisher Scientific). For real-time PCR, 1 µg of RNA was used to synthesize cDNA using the QuantiTect reverse transcription kit (Qiagen). qPCR was carried out on the Step-One Plus real-time PCR system (Applied Biosystems) using primaQUANT ADVANCED qPCR Master Mix (Steinbrenner). Transcript levels were calculated following the X0-method (19) and normalized to NUDC. The sequences of the PCR primers are provided in Supplementary Table S1.

### RNA sequencing and transcript expression analysis

For RNA sequencing, DNase-treated total RNA samples used for transcriptome analysis were quantified using the Qubit RNA HS Assay (Thermo Fisher Scientific), and RNA quality was assessed via capillary electrophoresis with the Fragment Analyzer using the ‘Total RNA Standard Sensitivity Assay’ (Agilent Technologies, Inc., Santa Clara, USA). All samples showed high RNA quality numbers (RQN; mean = 9.96). Library preparation followed the manufacturer’s protocol with the ‘VAHTS™ Stranded mRNA-Seq Library Prep Kit for Illumina®’. In short, 300 ng of total RNA was used for mRNA capture, fragmentation, cDNA synthesis, adapter ligation, and library amplification. The libraries were bead-purified, normalized, and sequenced on the HiSeq 3000/4000 system (Illumina Inc., San Diego, USA) with a single-read (SR) 1×150 bp setup. The bcl2fastq tool was used for file conversion, adapter trimming, and demultiplexing.

Data analysis on fastq files was conducted using CLC Genomics Workbench (version 12.0.3, QIAGEN, Venlo, NL). Reads were trimmed for Illumina TruSeq adapters and quality (default settings: bases below Q13 were trimmed from the ends, and a maximum of 2 ambiguous nucleotides were allowed). Reads were then mapped to the Mus musculus genome (mm10; GRCm38.86, March 24, 2017). Statistical analysis was performed with Qlucore Omics Explorer software. TPM values were log2-transformed and prefiltered (threshold 0.1). Depending on the experimental conditions, either multi-group or two-group comparisons were used. Genes with an absolute fold change of >1.5 and p<0.05 were considered differentially expressed (DGE). DGE data were further analyzed using Ingenuity Pathway Analysis (IPA) (Qiagen Inc., 2016) to identify potential upstream regulators contributing to the observed transcriptional changes.

### Western Blot

Hearts were homogenized using a Tissue Ruptor (Qiagen, Hilden) according to manufacturer’s protocol in RIPA buffer supplemented with protease and phosphatase inhibitors. Protein concentration was measured using a bicinchoninic acid (BCA) protein assay kit (Thermo Scientific). Equal amounts of protein were loaded onto a 7.5% or 10% SDS-PAGE gel, separated, and transferred onto Protran nitrocellulose membranes using the Pierce™ G2 Fast Blotter. Membranes were blocked with Odyssey blocking buffer (LI-COR Biosciences) and probed with antibodies against BDH1 (15417-1-AP), PDK4 (12949-1-AP), OXCT1/SCOT (12175-1-AP), all from Proteintech, CPT2 (PA5-19222, Thermo Fisher), PLIN2 (ab52356, Abcam), PLIN5 (GP31, Progen), and HMGCS2 (#20940, Cell Signaling). The secondary antibodies used were anti-rabbit IRDye800CW or anti-Rabbit IRDye680RD, anti-goat IRDye800CW, and anti-guinea pig IRDye680RD (LI-COR Biosciences). Signals were visualized and quantified using the Odyssey near-infrared imaging system (LI-COR Biosciences).

### Myocardial energy and substrate metabolism

Myocardial energy and substrate metabolism was analyzed by extracellular flux analysis in intact high-precision, vibratome-cut cardiac tissue slices (150 µm) using a Seahorse XFe24 analyzer (Agilent) based method that has been recently developed by our group (20).

### Mitochondrial isolation and respirometry

Cardiac mitochondria were isolated by differential centrifugation as described before (21) with moderate modifications. Briefly, ventricles were excised, placed in an isolation buffer [in mmol/l: 200 mannitol, 50 sucrose, 5 KH_2_PO_4_, 5 3-(*N*-morpholino)propanesulfonic acid (MOPS), and 1 EGTA, with 0.1% bovine serum albumin (BSA), pH 7.15 (adjusted with KOH)], and minced into small pieces. The suspension was homogenized using a Potter-Elvehjem-Homogenisator in isolation buffer containing 5 U/ml protease (four strokes, 800 rpm), and for another four strokes after addition of isolation buffer to dilute protease. The suspension was centrifuged at 3.200 *g* for 10 min; the pellet was resuspended in 5 ml of isolation buffer and centrifuged again at 500 *g* for 10 min. Next, the supernatant was centrifuged at 3.200 *g* for 10 min, and the final pellet was suspended in 0.4 ml of isolation buffer and kept on ice. The protein content was determined by the Bradford method (22). All isolation procedures were conducted at 4°C. Mitochondrial respiration (0.1 mg/ml protein concentration) was measured after administration of 5 mmol/l succinate in combination with 10 µmol/l rotenone (state 2), and after ADP (1 mmol/l) stimulation (state 3). Respiratory control ratios (RCR) were calculated as state 3 / state 2. In separate experiments, ketone body-related mitochondrial respiratory capacity was assessed by administration of 5 mmol/l β-hydroxybutyrate (Biomol, AG-CR1-3616) to ADP stimulated mitochondria.

### Statistical Analysis

The experiments were performed in a randomized and blinded fashion. Data are reported as mean ± SD. Statistical analyses, excluding RNAseq data, were conducted using GraphPad Prism 10. All results were tested by either two-tailed unpaired t test or one-way ANOVA followed by post hoc test as indicated. A *p-* value of less than 0.05 was considered statistically significant.

## Results

### DIO-STZ mice show a hyperglycemic, hypoinsulinemic diabetic phenotype

All mice underwent the diet feeding protocol beginning at the age of 9 weeks (Figure 1A). Starting at the 3rd week of feeding, both groups receiving the calorie rich HFHSD, i.e. the DIO-STZ and the DIO group, showed increased body weights compared to the two groups that were fed a standard chow, i.e. the chow and the STZ group (Figure 1B). After 5 weeks of feeding, animals of the DIO-STZ and the STZ groups received low-dose STZ injections to induce partial destruction of pancreatic β-cells. In the case of the DIO-STZ group, the body weight decreased to the level observed in the control animals within one week after STZ treatment (Figure 1B). At the end of the protocol, i.e. after 11 weeks, the body weight of DIO-STZ animals was comparable to chow group (31.5±2.0 g vs. 31.2±2.3 g), whilst DIO animals had increased body weight (42.2±4.4 g), and STZ animals showed lower weight (28.3±2.0 g) (Figure 1B).

**Figure 1:**
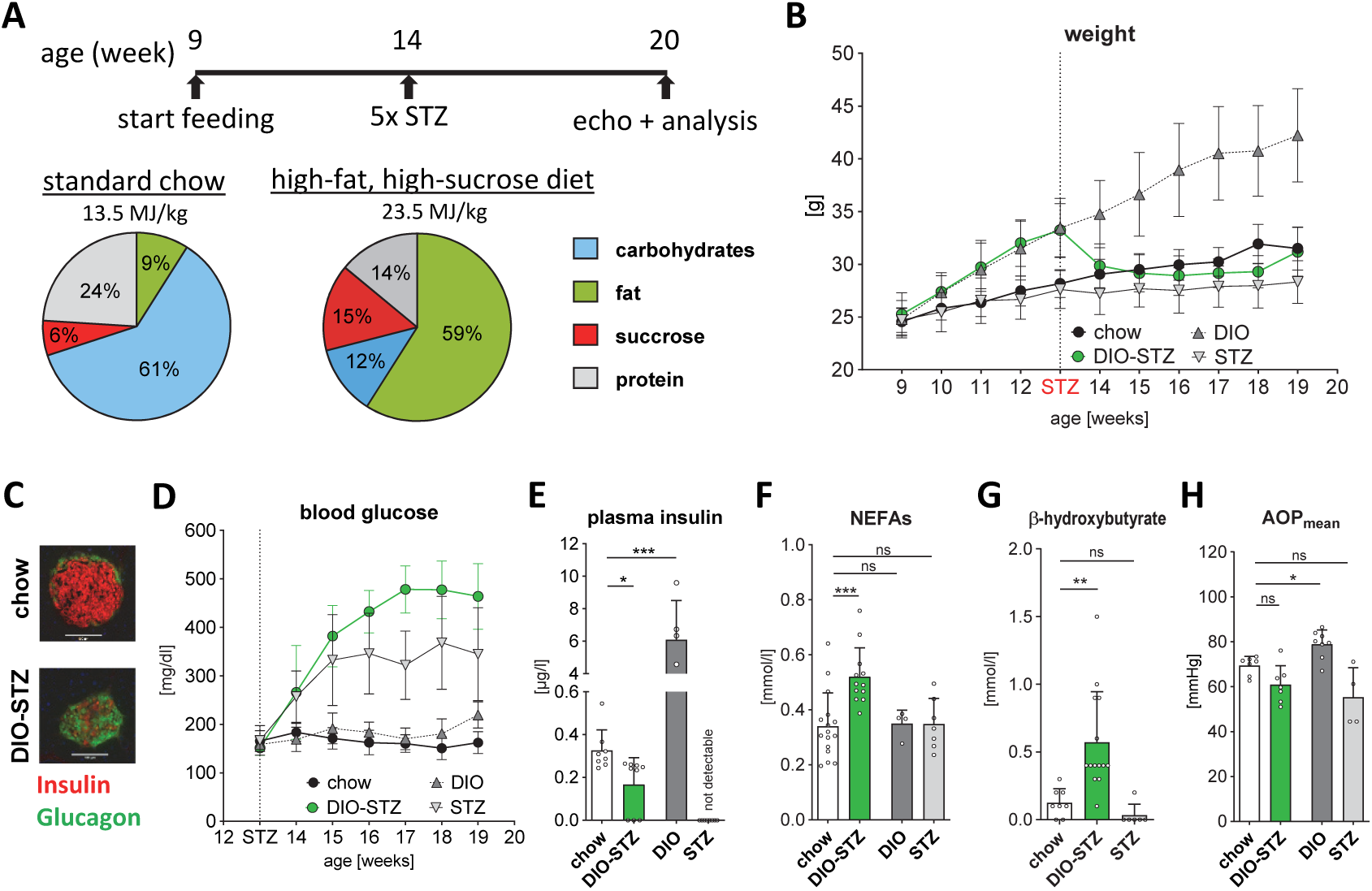
DIO-STZ treatment protocol and general characterization of the model. (A) Upper panel: For induction of T2DM, male C57BL/6J mice were fed a high-fat high-sucrose diet (HFHSD) over a period of 11 weeks in combination with additional injections of streptozocin in the 5th week (DIO-STZ group). Control animals received standard chow and no STZ injections (chow group). As additional groups, the DIO group were fed a HFHSD without STZ injections, and animals of the STZ group received STZ, and were fed standard chow. Lower panel: Energy composition of the standard chow and the HFHSD. (B) Summarized data for body weight development over the experimental protocol (n=12-32). (C) Representative immunofluorescence images of the STZ destroyed beta cell mass in the Langerhans islets (left), and summarized data of plasma blood glucose levels starting from the time point of STZ injections (n=8-16). (D-F) Summarized data of plasma levels of insulin (n=5-9), non-esterified fatty acids (n=4-16), and β-hydroxybutyrate (n=6-14). (G) Mean aortic pressure (AOP_mean_; n=4-8). Data are shown as means ± SD. Statistical analysis was performed using one-way ANOVA followed by Dunnett’s *post hoc* test with chow as control. \**P* < 0.05, \*\**P* < 0.01, \*\*\**P* < 0.001, ns = not significant.

Following the administration of STZ, a significant increase in blood glucose levels was observed in both the DIO-STZ and the STZ groups with higher glucose levels in the DIO-STZ mice compared to STZ (478± 8 mg/dl vs. 368±95 mg/dl) starting from the 2nd week after STZ injection (Figure 1D). These two groups also showed reduced plasma insulin levels as result of the partial β-cell destruction (Figure 1E), which is also supported by fluorescence microscopy of the islets of Langerhans, which showed a clearly degenerated beta cell mass (Figure 1C). HFHSD feeding alone did not induce an increase in blood glucose levels (Figure 1D), but resulted in a pronounced increase in plasma insulin levels (Figure 1E) pointing towards a reduced insulin sensitivity. The unique metabolic phenotype of the DIO-STZ model is also underlined by the observation that only DIO-STZ mice showed a robust increase in plasma free fatty acid levels compared not only to chow, but also to STZ mice (Figure 1F). In addition, DIO-STZ mice showed elevated beta-hydroxybutyrate plasma levels, (Figure 1G), and were normotensive (Figure 1H).

Erythrocyte count, hemoglobin concentration, and hematocrit levels were slightly lower in DIO-STZ compared to chow animals, and no differences in red blood cell indices, white blood cell count, and platelet count were found (Supplementary Table S2).

### DIO-STZ hearts develop a HFpEF phenotype

To investigate whether the diabetes and obesity related alterations in whole body metabolism affect cardiac function, high-resolution echocardiography was performed at experimental week 11 (Figure 2A). Here, the male DIO-STZ T2DM mice showed a pronounced reduction in cardiac output by more than 40 % accompanied by a completely preserved ejection fraction, and a robust reduction in the EDV. Interestingly, the end-diastolic pressure in the DIO-STZ group was increased compared to non-diabetic chow group (Figure 2B) despite reduced EDV indicating diastolic dysfunction with increased filling pressures. In summary, male DIO-STZ mice present a HFpEF phenotype, which is also confirmed by speckle tracking based strain analysis. Here, a reduction in global longitudinal strain is detected in DIO-STZ mice (Figure 2C). In addition, the occurrence of a diastolic dysfunction is supported by a reduced diastolic (reverse peak) global longitudinal strain rate (Figure 2C). Finally, also an increased relative wall thickness (RWT) further substantiated the presence of a HFpEF phenotype in the DIO-STZ model (Figure 2D). Please note that the increase in RWT was not the consequence of myocardial hypertrophy as DIO-STZ animals showed a slight reduction in both heart weights (Figure 2E) and heart-to-body weight ratios (Figure 2F) compared to chow controls.

**Figure 2:**
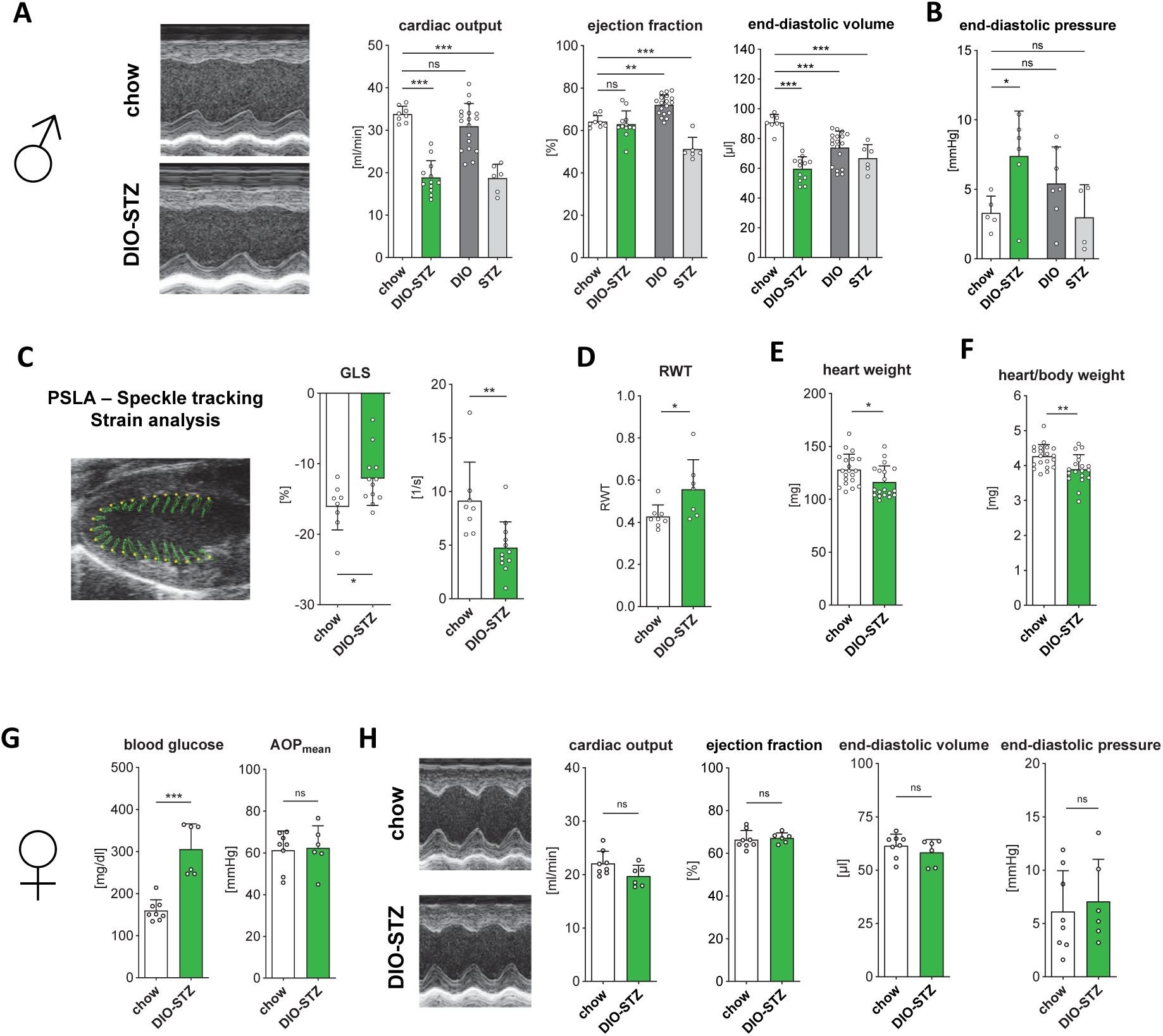
Cardiac phenotyping *in vivo*. (A) Representative echocardiographic SAX m-mode images of male chow and DIO-STZ hearts (left), and summarized data for cardiac output, ejection fraction, and end-diastolic volume (n=6-18). (B) Summarized data of end-diastolic pressure obtained by left ventricular catheterization (n=4-7). (C) Example image of speckle-tracking based left-ventricular strain analysis on echocardiographic recording in parasternal long axis view (PSLA). Green lines indicate endocardial displacement over a complete heart cycle (left). Summarized data for global longitudinal strain (GLS), and reverse peak (diastolic) global longitudinal strain rate (GLSR)(n=8-12). (D) Summarized data for relative wall thickness (RWT) (n=7-8), (E) heart weight and (F) heart to body weight ratio. (G) Phenotyping of female DIO-STZ animals. Left: Summarized data of plasma blood glucose levels and mean aortic pressure (AOP_mean_; n=6-8). Right: Representative echocardiographic SAX m-mode images of female chow and DIO-STZ hearts, and summarized data for cardiac output, ejection fraction, end-diastolic volume, and end-diastolic pressure obtained by left ventricular catheterization (n=6-8). Data are shown as means ± SD. Statistical analysis was performed using one-way ANOVA followed by Dunnett’s *post hoc* test with chow as control (A-B), or by unpaired *t*-test (C-F). \**P* < 0.05, \*\**P* < 0.01, \*\*\**P* < 0.001, ns = not significant.

Interestingly, the HFpEF development is clearly a unique characteristic of the two-hit DIO-STZ model as the HFHSD fed DIO group showed no signs of cardiac dysfunction or altered ventricular filling, and the STZ group showed a reduced ejection fraction without indication of increased ventricular filling pressure (Figure 2A & B).

Additional echocardiographic analysis of cardiac function demonstrated that the cardiac HFpEF phenotype was already developed at an earlier time point of 9 weeks of HFHSD, i.e. 4 weeks after STZ treatment (Supplementary figure 1). Here, cardiac function of DIO-STZ hearts was characterized by a 23% reduction in cardiac output, a preserved ejection fraction (58.3±3.1% vs. 57.6±4.1%), and a reduction in end-diastolic volume.

In addition, to investigate potential sex-dependent differences in vulnerability to HFpEF development, female C57Bl/6J mice underwent the experimental DIO-STZ protocol described above. Here, DIO-STZ treatment for 11 weeks resulted in a robust increase in blood glucose, a slightly reduced body weight and increased beta-hydroxybutyrate plasma levels compared to chow mice, and no difference in blood pressure was detected (Figure 2G, and Supplementary Figure S2). Cardiac phenotyping of female mice revealed no indication of HFpEF development as seen by no differences in cardiac output, ejection fraction, ventricular volumes, and end-diastolic pressure (Figure 2H, and Supplementary Figure S2).

Furthermore, there is evidence of varying susceptibility to HFpEF development between the C57Bl/6 substrains C57Bl/6J and C57Bl/6N, and this difference in HFpEF development could be clearly attributed to the presence of a loss-of-function mutation in the nicotinamide nucleotide transhydrogenase (*Nnt*) gene (23). In this line, we performed an additional experimental series with male C57Bl/6N mice. First, the verified the presence of the *Nnt* mutation in the C57Bl/6J substrain, and the absence of the mutation in the C57Bl/6N substrain by PCR analysis (Supplementary Figure S3). At the end of the experimental diabetes protocol, DIO-STZ mice showed elevated blood glucose and beta-hydroxybutyrate levels, whilst body weight was slightly reduced and blood pressure was unchanged (Supplementary Figure S4A & B). Cardiac phenotyping revealed a strong reduction in cardiac output whilst ejection fraction was not altered in DIO-STZ animals pointing towards an HFpEF phenotype (Supplementary Figure S4C). This assumption was supported by reduced left-ventricular volume accompanied by increased end-diastolic pressure (Supplementary Figure S4C & B).

### Intrinsic disturbances of contractile myocardial function and vascular regulation

We performed an extensive cardiac phenotyping in the isolated perfused Langendorff heart to further characterize whether the observed cardiac dysfunction described above is a consequence of intrinsic myocardial deteriorations, and not caused by extrinsic factors, e.g. altered influence of the vegetative nervous system or altered substrate supply to the heart.

First, to analyze myocardial compliance, we increased stepwise the volume of the left ventricular balloon and determined the resulting increase in LVP_min_ (Figure 3A). Here, no difference in increase of LVP_min_ was observed during the first 5 to 6 gradual volume increases. Notably, these volume injections resulted in a physiologically relevant increase in ventricular preload as seen by a Frank-Starling-Mechanism mediated increase in LVP_max_ (Figure 3A-B, marked area). During the following volume steps, DIO-STZ hearts showed a trend towards a less steep increase in LVP_min_ indicating that the intrinsic ventricular compliance is not increased compared to chow hearts. In addition, as myocardial remodeling with increased tissue fibrosis would also affect myocardial tissue stiffness, we performed histochemical Sirius red stainings from chow and DIO-STZ hearts (Figure 3C). The collagen tissue content determined by polarized light image analysis was not different between chow and DIO-STZ animals (Figure 3D).

**Figure 3:**
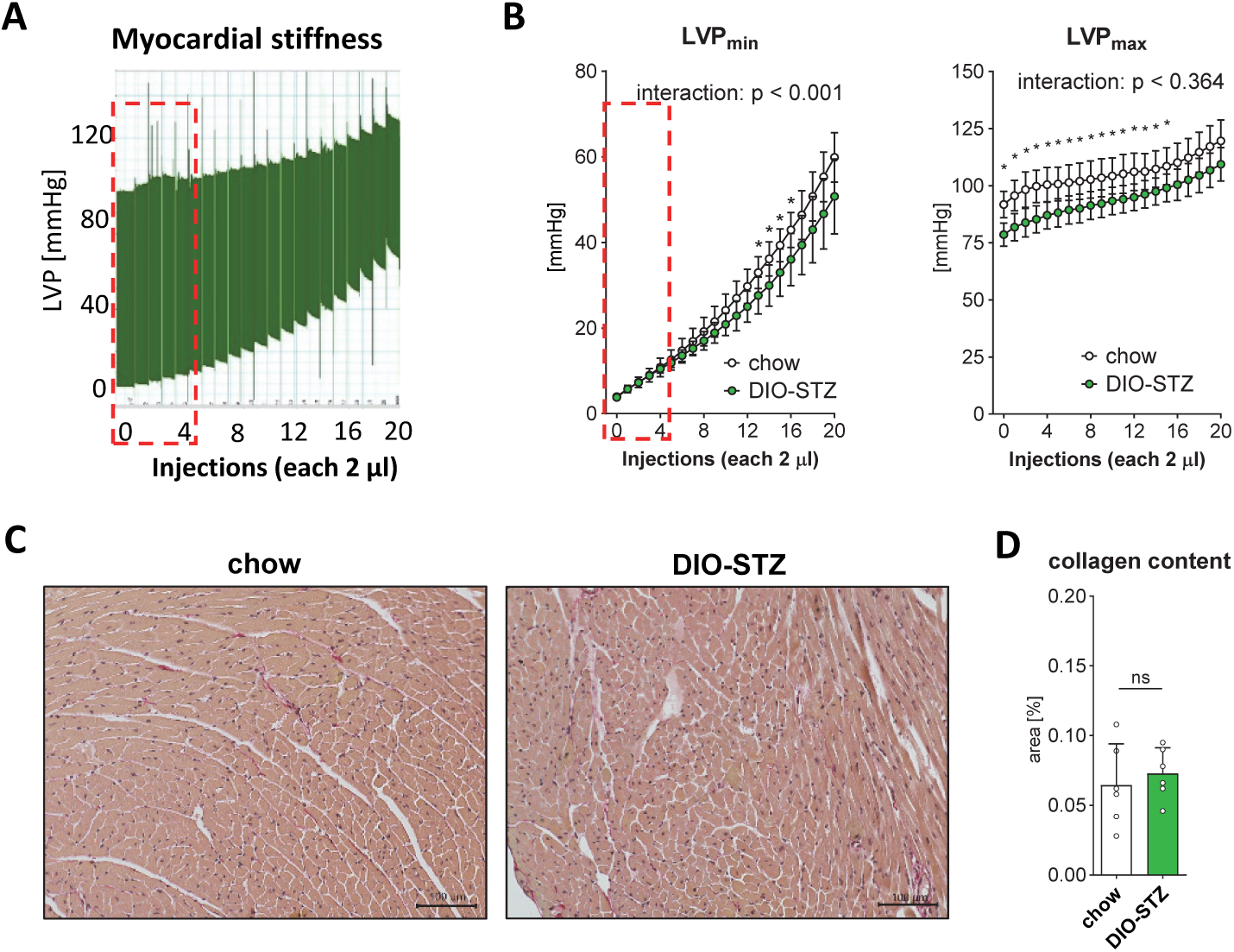
Intrinsic myocardial stiffness, and myocardial collagen content. (A) Example trace of determination of volume-pressure relationship: Effect of increased ventricular preload by stepwise increase in ventricular volume (2 µl per injection). The area marked with a red dotted line indicates the presumed physiological range of preload increases, as a Frank-Starling-mechanism induced effect on developed pressure can be seen here. (B) Summarized data for effect on preload increase on LVP_min_ (left), and LVP_max_ (right)(n=6 per group). (C) Histological example images of Sirius red stained ventricular tissue (D) Summarized data of collagen content analysis (n=6 hearts per group). Data are shown as means ± SD. Statistical analysis was performed using two-way ANOVA followed by Tukey’s *post hoc* test (B), or by unpaired *t*-test (D). \**P* < 0.05, ns = not significant.

Second, the intrinsic myocardial pump function was investigated. Systolic isovolumetric contraction work after applying comparable preload (LVP_min_) was lower in DIO-STZ hearts as seen by a reduction in LVP_max_ of 16% (76.1±5.2 mmHg vs 90.1±6.9 mmHg)(Figure 4A). Furthermore, no difference in maximum speed of left ventricular pressure rise (dP/dt_max_) was observed, whereas maximum speed of pressure decay (dP/dt_min_) was reduced in DIO-STZ hearts potentially pointing towards an additional diastolic component of the cardiac dysfunction.

**Figure 4:**
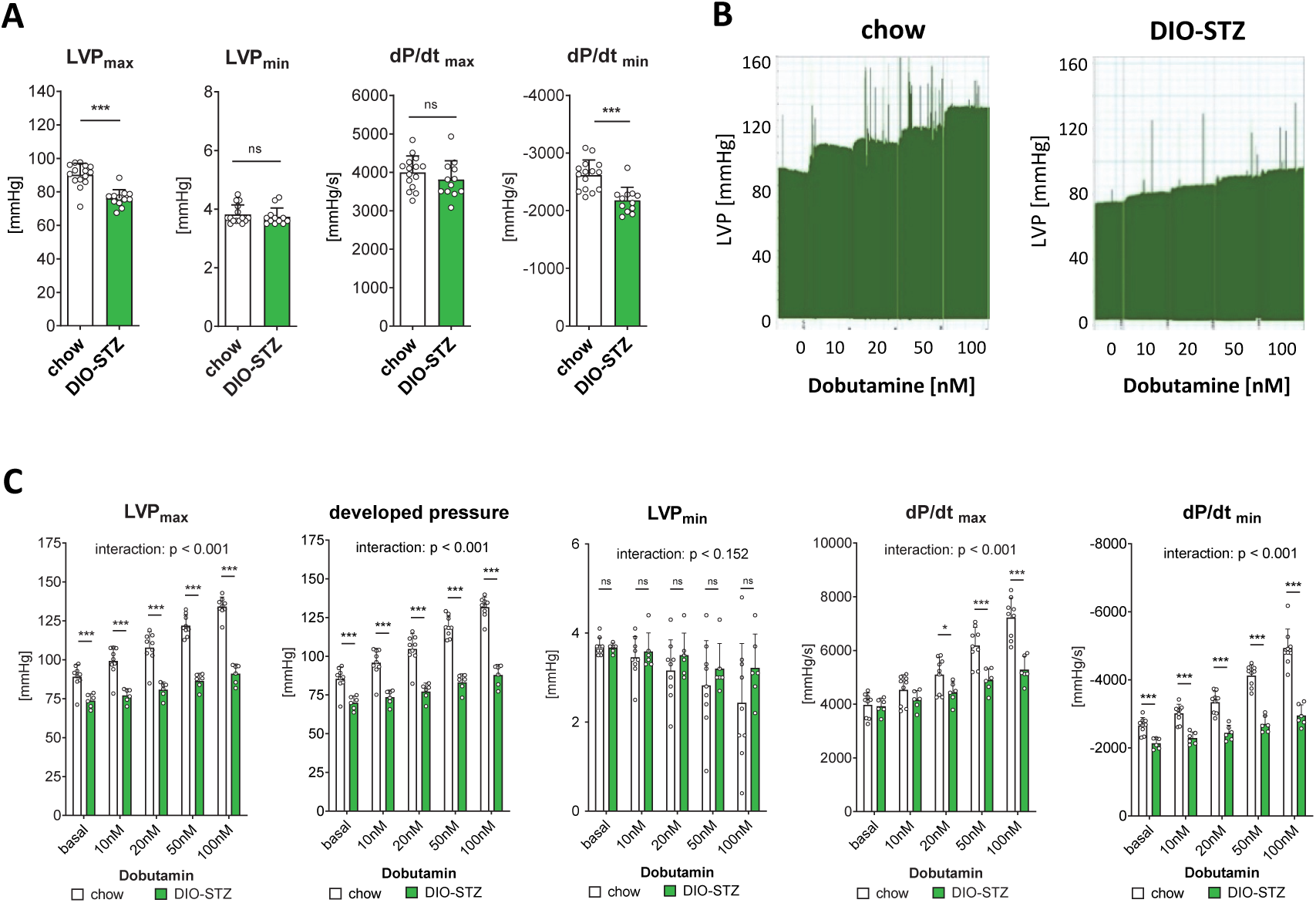
Characterization of contractile function, and dobutamine stress test in the isolated perfused heart. (A) Contractile function under basal conditions: Summarized data for LVP_max_, LVP_min_, dP/dt_max_, and dP/dt_min_ of DIO-STZ or chow fed control isolated hearts (n=12-15 per group). (B) Example traces of ventricular pressure response to β-adrenergic stimulation by dobutamine. (C) Summarized data for dobutamine effects on LVP_max_, developed pressure, LVP_min_, dP/dt_max_, and dP/dt_min_ of DIO-STZ or chow fed control isolated hearts (n=6-9 per group). Data are shown as means ± SD. Statistical analysis was performed using unpaired *t*-test (A), or two-way ANOVA followed by Tukey’s *post hoc* test (C). \**P* < 0.05, \*\**P* < 0.01, \*\*\**P* < 0.001, ns = not significant.

Third, as HFpEF patients often only show symptoms under stress conditions, we tested the cardiac response to β-adrenergic stimulation with dobutamine (Figure 4B-C). In chow hearts, dobutamine treatment resulted in a concentration dependent increase in LVP_max_, developed pressure, dP/dt_max_ and dP/dt_min_, and a decrease in LVP_min_, respectively. All these dobutamine effects were strongly blunted or not detectable in DIO-STZ hearts.

### DIO-STZ hearts show alterations in key metabolic enzymes

Principal component analysis and hierarchical cluster analysis of differentially regulated genes revealed that the gene expression profile of DIO-STZ mouse hearts formed a distinct cluster separating not only from chow-fed animals but also from both DIO and STZ groups (Figure 5A). Ingenuity pathway analysis further demonstrated that three of the top four most significantly regulated canonical pathways were linked to lipid and fatty acid metabolism, including “Mitochondrial Fatty Acid Beta-Oxidation,” “Regulation of Lipid Metabolism by PPARalpha,” and “Fatty Acid β-Oxidation” (Figure 5B). qPCR data collected from additional independent chow-fed and DIO-STZ mouse hearts supported the increased expression of *Cpt1b*, *Cpt2* and *Pdk4* genes and thus confirmed alterations in myocardial substrate metabolism (Figure 5C). In line with the RNA data, altered expression of proteins related to lipid and fatty acid metabolism were detected, e.g. increase in CPT2, PLIN2, PLIN5 and PDK4 supporting an elevated fatty acid metabolism and suppression of glucose utilization. Moreover, decrease in BDH1 and OXCT1/SCOT expression as well as increase in HMGCS2 expression were found in DIO-STZ hearts, pointing towards alterations in ketone body metabolism (Figure 5D-E).

**Figure 5:**
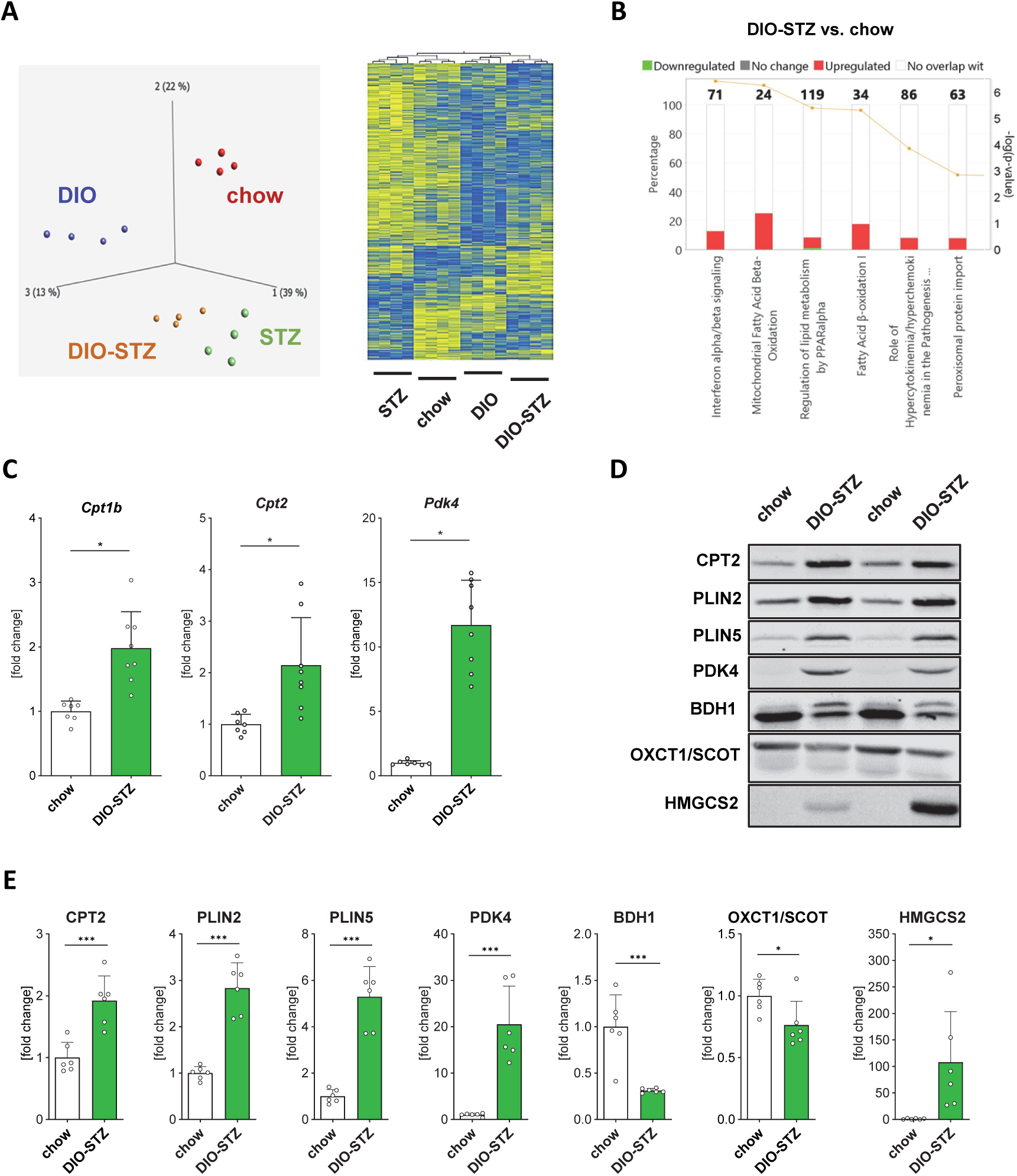
DIO-STZ hearts show alterations in key metabolic enzymes. (A) Myocardial bulk RNA-Seq analysis was performed in DIO-STZ and chow mice, as well as in both STZ and DIO mice. Principal component analysis as well as hierarchical cluster analysis of differentially regulated genes. (B) Ingenuity pathway analysis revealed that three of the four most strongly regulated canonical pathways in DIO-STZ hearts are related to lipid and fatty acid metabolism. (C) Summarized qPCR data for RNA expression of key metabolic enzymes *Cpt1b*, *Cpt2* and *Pdk4* (n=7-8). (D-E) Example western blots, and summarized data of CPT2, PLIN2, PLIN5, PDK4, BDH1, OXCT1/SCOT, and HMGCS2 protein levels (n=6 per group). Data are shown as means ± SD. Statistical analysis was performed using unpaired t-test. \**P* < 0.05, \*\**P* < 0.01, \*\*\**P* < 0.001, ns = not significant.

### DIO-STZ hearts show improved capacity to oxidize fatty acids

To investigate the functional relevance of the DIO-STZ related alterations in gene and protein expression, we performed extracellular flux analysis on intact cardiac tissue slices, a method that was recently developed by our group (20)(Figure 6A). Myocardial tissue from DIO-STZ animals showed increased oxygen consumption rates (OCR) under resting conditions (OCR basal) as well as after stimulation of respiration by mitochondrial uncoupling (OCR uncoupled) compared to the chow group when a substrate combination of glucose and palmitate was offered (figure 6B). Furthermore, inhibition of either CPT1 by etomoxir or MPC by UK5099 to block fatty acid or glucose oxidation, respectively, revealed elevated fatty acid oxidation capacity in the DIO-STZ group, and in contrast, no effect on glucose oxidation capacity was detected. To support the finding that fatty acid oxidation capacity is increased and carbohydrate oxidation capacity is not affected, we performed additional experiments with either a substrate combination of glucose and pyruvate, or palmitate alone (Figure 6C). Here, no difference in uncoupled OCR was found between cardiac tissue from DIO-STZ and chow animals with glucose and pyruvate as sole substrates (Figure 6D), whilst uncoupled OCR was two-fold increased in DIO-STZ when solely palmitate was offered further supporting the finding of increased myocardial capacity to oxidize fatty acids (Figure 6E).

**Figure 6:**
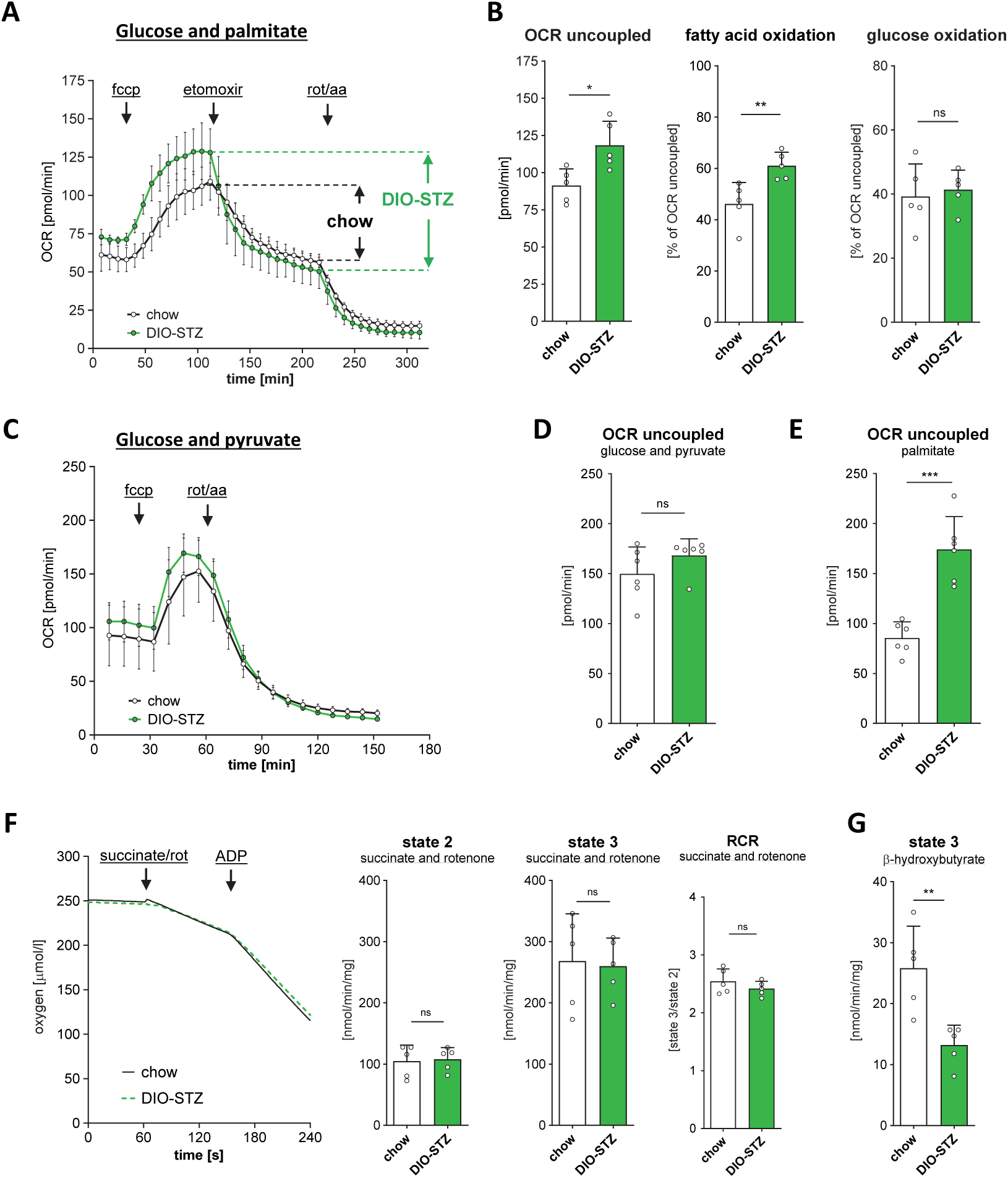
Analysis of myocardial energy and substrate metabolism. (A) Summarized traces of extracellular flux analysis in intact cardiac tissue slices of chow and DIO-STZ animals (n=5 per group) using glucose and palmitate as substrates. Oxygen consumption rates (OCR) were determined under resting conditions (basal), and after mitochondrial uncoupling using carbonyl cyanide-p-trifluoromethoxyphenylhydrazone (fccp). The oxidation capacity for fatty acids was investigated by inhibition of carnitine palmitoyltransferase I using etomoxir, and for glucose by inhibition of mitochondrial pyruvate carrier using UK5099, respectively. The shown traces represent experiments with etomoxir; experiments with UK5099 were performed in parallel. (B) Summarized data of OCR uncoupled as well as summarized data of FFA and glucose oxidation capacity, respectively. (C) Summarized traces of extracellular flux analysis in intact cardiac tissue slices of chow and DIO-STZ animals (n=6 per group) using glucose and pyruvate as substrates. Oxygen consumption rates (OCR) were determined after mitochondrial uncoupling using fccp. The shown traces represent experiments with glucose and pyruvate; experiments with palmitate as sole substrate were performed in parallel. (D-E) Summarized data of OCR uncoupled in experiments with either (D) glucose and pyruvate or (E) palmitate. (F) Summarized traces (left) and data (right) of mitochondrial respiration experiments in isolated heart mitochondria from chow and DIO-STZ animals (n=5 per group). Mitochondrial state 2 respiration was initiated by addition of succinate (in combination with rotenone), and state 3 respiration was initiated by addition of ADP. Respiratory control ratio was calculated as state 3 / state 2. (G) Summarized data of state 3 respiration in β-hydroxybutyrate supported isolated mitochondria from chow and DIO-STZ animals (n=5 per group). Data are shown as means ± SD. Statistical analysis was performed using unpaired t-test. \**P* < 0.05, \*\**P* < 0.01, \*\*\**P* < 0.001, ns = not significant.

In addition to glucose and fatty acids, which contribute the most to total myocardial ATP production, the heart is also able to metabolize other substrates including ketone bodies, especially under pathophysiological conditions. To investigate the impact of the reduced expression of BDH1 and OXCT1/SCOT in DIO-STZ hearts, we performed respirometric measurements in isolated heart mitochondria. Using succinate as substrate no difference was found in state 2 and state 3 respiration as well as in the RCR pointing towards normal function of the mitochondrial electron transport chain of DIO-STZ hearts (Figure 6F). In contrast, the mitochondrial capacity to oxidize the ketone body beta-hydroxybutyrate was about 50% lower in the DIO-STZ group than in the chow group (Figure 6G).

## Discussion

Here, we show that the DIO-STZ mouse model mirrors metabolic features of T2DM including hyperglycemia, hypoinsulinemia, elevated fatty acid and ketone body levels. Notably, DIO-STZ mice develop a robust HFpEF phenotype, which is mainly characterized by a diastolic dysfunction with increased filling pressures. The development of HFpEF is unique for the two-hit DIO-STZ model as HFpEF was neither found in mice that received only STZ-treatment nor in mice that were solely fed a HFHSD. Importantly, the systemic, diabetes-related metabolic alterations of the DIO-STZ model resulted in myocardial dysfunction in the isolated heart pointing towards secondary intrinsic disturbances of the myocardium. Central differences between the DIO-STZ model, which develops HFpEF on the one hand, and the STZ or the DIO models, which do not develop such an HFpEF phenotype, on the other, lie in specific changes in metabolic parameters. Our data suggest that changes in glucose and insulin homeostasis alone are not sufficient to induce a HFpEF phenotype in the observed period, as neither hypoinsulinaemic hyperglycaemia as in the STZ model nor normoglycaemic hyperinsulinaemia as in the DIO model induce such cardiac dysfunction. Interestingly, two metabolic changes that are only altered in the DIO-STZ model are related to fat metabolism, namely increased plasma levels of FFAs and the ketone body β-hydroxybutyrate, which mainly results from hepatic fatty acid breakdown. This link to alterations in lipid metabolism might be of relevance as obesity is now seen as a key player in HFpEF development (24, 25).

Although there is clear evidence that obesity and diabetes are strongly associated with the occurrence of heart failure, the causal relationship is incompletely understood. A potential factor contributing to this knowledge gap is the limited availability of suitable mouse models for investigating the molecular and systemic relationships in HFpEF.

In general, HFpEF is a multifactorial syndrome in which the triggering factors and pathophysiological processes can vary considerably, and it is therefore unlikely that a single mouse model can reflect this mechanistic variability in the sense of ‘one size fits all’. Several murine models for HFpEF have been described and their benefits and shortcomings have been discussed in the recent years (26–30). Here, a large group of these models are related to common HFpEF comorbidities as hypertension and/or obesity and diabetes (genetic as well as non-genetic). The DIO-STZ model was first described in 2000 by Reed et al as a new T2DM rat model for the testing of antidiabetic compounds (13). There is some evidence both from rats and mice that high-fat diet and streptozotocin treatment can initiate a diastolic dysfunction or HFpEF phenotype (31–33). Here, a critical role for SIRT6 mediated alterations in endothelial lipid transport from the blood towards the tissue cardiac (33).

In the present study, we focused less on the significance of the endothelium within the DIO-STZ HFpEF mouse model than on extensive functional and metabolic phenotyping of the myocardium. Our analysis of the DIO-STZ model revealed a pronounced cardiac manifestation of the T2DM in the sense of a fully developed HFpEF phenotype with increased LV filling pressures and RWT, and reduced cardiac output, end-diastolic volume, GLS, and GLSR. In addition to this, bulk-sequencing analysis of myocardial RNA from DIO-STZ mice revealed extensive changes in the expression of genes related to substrate metabolism. Genes involved in fatty acid metabolism (e.g. *Cpt1b* or *Cpt2),* regulation of metabolic flexibility (e.g. *Pdk4*), or ketone body metabolism (*Bdh1, Oxct1* or *Hmgcs2*) in particular showed altered expression. The functional relevance of this altered gene expression is supported by a highly increased myocardial tissue capacity to oxidize palmitate, a finding that is in line with recent observations that the contribution of fatty acid oxidation to overall ATP production is increased in cardiometabolic HFpEF (34, 35).

Next to this regulation of fatty acid metabolism, there is increasing evidence that ketone bodies act as relevant substrate for the heart under physiological as well as under pathophysiological conditions including HFpEF. Although the literature is not consistent at this point, recent evidence indicated an impairment of myocardial ketone oxidation in HFpEF patients as well as in mice with cardiometabolic HFpEF (35, 36). Interestingly, we also found a robust reduction in myocardial BDH1 and SCOT expression. In addition, the mitochondrial capacity to oxidize βOHB in the DIO-STZ model was attenuated demonstrating that the molecular alterations translated into a functional depression. Our data are in line with recent findings in the combinatory ‘two-hit’ model of high fat diet and nitric oxide synthase inhibitor L-NAME treatment (HFD+L-NAME), one of the best described HFpEF models (37). In this model, Sun et al. recently also found a reduced myocardial expression of the ketolytic enzymes BDH1 and SCOT and, concomitantly, a reduced rate of βOHB oxidation in the isolated working heart (35). Thus, although there are clear differences in the HFpEF etiology between the hypertensive HFD-L-NAME model, and the normotensive DIO-STZ model with a fully developed T2DM-like diabetic phenotype, a compromised ketone metabolism appears to be a common feature of metabolically linked HFpEF development. Although the comparable changes in ketone body metabolism are striking, their role as HFpEF driving factor is questionable because improving myocardial ketone oxidation rates experimentally did not result in improvement of cardiac dysfunction in HFD+L-NAME mice (35). However, it remains unclear whether the altered ketone body metabolism is causal or merely correlative to HFpEF development in in the DIO-STZ model.

The C57Bl/6J substrain represents a frequently used genetic background strain in transgenic mice, and the DIO-STZ model leads to robust HFpEF phenotype also in this mouse strain. This contrasts in particular with the HFD+L-NAME model, in which mice of the C57BL/6J substrain show no or only mild HFpEF development (23). Mechanistically, Pepin et al. described that the development of a HFpEF phenotype in the HFD+L-NAME mouse model depends on the presence of a functional nicotinamide nucleotide transhydrogenase (NNT) (23), which is inactivated due to a 5-exon deletion (exons 7-11) in the 6J but not 6N strain. A key physiological role of the NNT is to maintain mitochondrial redox balance by restoration of the NADPH pool at the expense of NADH oxidation, thereby linking energy metabolism with antioxidative defense. Importantly, our present study demonstrated that DIO-STZ treatment initiated a HFpEF phenotype occurring in both the C57Bl/6J and the C57Bl6N substrain, clearly pointing towards divergent underlying molecular disease mechanisms between the cardiometabolic HFD+L-NAME model and the diabetic DIO-STZ model.

Mechanistic differences between preclinical HFpEF mouse models do not only exist with regard to the significance of the NNT, but also to other aspects of the disease, such as the emergence of mitochondrial changes with defects in oxidative phosphorylation and the respiratory chain or in the development of cardiac fibrosis. In theory, the occurrence of cardiac fibrosis would be a plausible explanation for an increase in myocardial stiffness. However, several findings argue against morphological tissue alterations as reason for the HFpEF phenotype in DIO-STZ mice. First, no difference in myocardial collagen content was observed indicating that the DIOSTZ mice did not develop cardiac fibrosis. Second, the marked increase in LV filling pressures with reduced end-diastolic volume occurring *in vivo* were not detectable *in vitro* in the isolated perfused heart, because no increase in volume-pressure relationship was detectable. If a morphological tissue remodelling with increased fibrosis resulting would cause the increased LV filling pressures, this structural remodeling, should also result in an increased stiffness in the isolated heart preparation, which, however was not measured. We interpret these results as evidence of the occurrence of a ‘functional stiffness’ caused by unknown factors in the *in vivo* situation, which, unlike ‘morphological stiffness’ caused by cardiac fibrosis, for example in the HFD-L-NAME model, is not detectable *in vitro*. To what extent changes in Ca^2+^ handling, titin and or troponin modifications may be involved remains to be investigated.

Epidemiological data suggest that women have at least a comparable or even an increased lifetime risk of developing HFpEF compared to men (38–40). However, the preclinical DIO-STZ protocol did not result in HFpEF development in female mice in contrast to male mice. This is in line with most preclinical mouse models showing that females develop no or only mild signs of HFpEF, and this protective effect seems to be independent of female sex hormones (23, 41, 42). Interestingly, the DIO-STZ treatment in female mice did not lead to HFpEF development, although a hyperglycemic, normotensive phenotype with elevated plasma ketone body levels was detected indicating that female mice were protected from T2DM related HFpEF development. Furthermore, this suggests that the mechanism by which female mice are protected from HFpEF development appear to be a more general mechanism as occurring in different HFpEF models with divergent etiologies, and therefore, unraveling the causative factors of these sex-specific differences in preclinical mouse models will be of relevance for a broader mechanistic understanding of HFpEF pathophysiology and may even help to identify possible molecular targets for HFpEF treatment.

## List of abbreviations

HFpEF: Heart failure with preserved ejection fraction
HFrEF: Heart failure with reduced ejection fraction
T2DM: Type 2 diabetes mellitus
SGLT2i: Sodium-glucose transport inhibitor
GLP-1RA: Glucagon-like peptide-1 receptor agonist
HFHSD: High-fat and high-succrose diet
HFD: High-fat diet
DIO: Diet-induced obesity
STZ: Streptozotocin
NEFA: Non-esterified fatty acid
ECG: Electrocardiogram
PSLAX: Parasternal long axis
SAX: Short axis
EDV: End-diastolic volume
ESV: End-systolic volume
LVPW: Left ventricular posterior wall
LVID: Left ventricular internal diameter
RWT: Relative wall thickness
GLS: Global longitudinal strain
GLSR: Global longitudinal strain rate
LV: Left ventricle
AOP_mean_: Mean aortic pressure
LVEDP: Left ventricular end-diastolic pressure
LVP_min_: Minimal left ventricular pressure
LVP_max_: Maximal left ventricular pressure
DGE: Differentially expressed gene
BDH1: Beta-hydroxybutyrate dehydrogenase
PDK4: Pyruvate dehydrogenase kinase 4
OXCT1: 3-oxoacid CoA-transferase 1
SCOT: Succinyl-CoA:3-ketoacid-CoA transferase
CPT1: Carnitine palmitoyltransferase 1
CPT2: Carnitine palmitoyltransferase 2
PLIN2: Perilipin-2
PLIN5: Perilipin-5
HMGCS2: 3-hydroxy-3-methylglutaryl-CoA synthase 2
RCR: Respiratory contral ratio
NNT: Nicotinamide nucleotide transhydrogenase
dP/dt_max_: Maximal speed of left ventricular pressure rise
dP/dt_min_: Maximal speed of left ventricular pressure decay
OCR: Oxygen consumption rate

## Declarations

### Authors’ contribution

AH, AS, SG, and AG conceived and designed research. AH, AS, ZF, LB, and KB performed experiments. AH, AS, SG, ZF, LB, and PP analyzed data. AH, AS, SG, KB, JF, and AG interpreted results of experiments. AH, AS, SG and AG prepared figures. AH drafted manuscript. SG and AG edited and revised manuscript. AH, AS, SG, ZF, LB, KB, JF, PP, and AG approved final version of manuscript.

## Acknowledgements

We would like to thank Julia Albrecht, Sevgi Bongartz, and Daniela Müller for their excellent assistance in conduction experiments. Furthermore, we would like to thank Drs. Altschmied and Haendeler for providing the PCR protocol and primers for genotyping the Nnt-mutation.

---

**Supplementary Figure 1:**
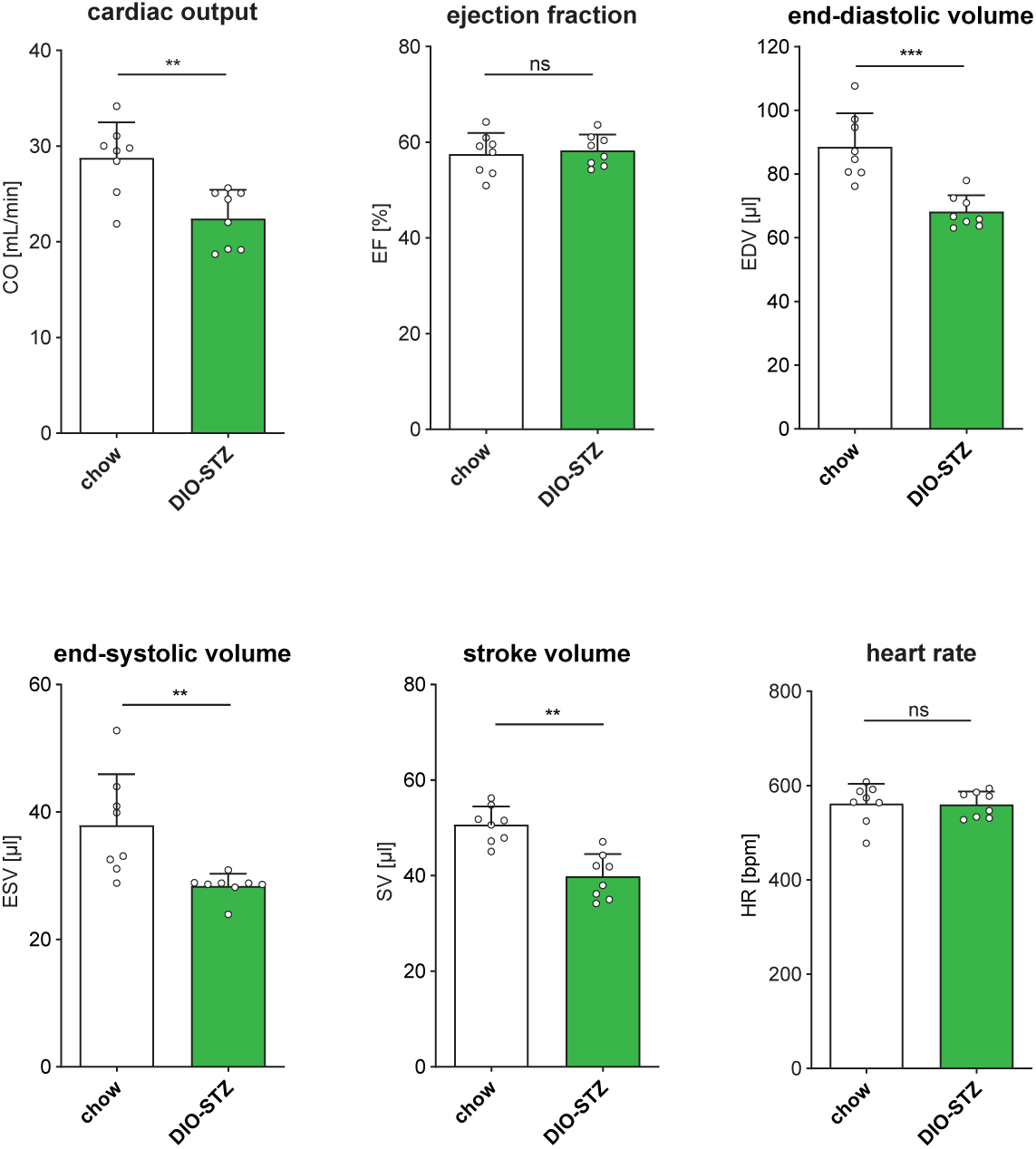
Cardiac functional phenotyping after 9 weeks of feeding. Summarized data for cardiac output, ejection fraction, and end-diastolic volume, end-systolic volume, stroke volume and heart rate in DIOO-STZ and chow animals after 9 weeks of feeding, i.e. 4 weeks after STZ (in the DIO-STZ group) (n=8). Data are shown as means ± SD. Statistical analysis was performed using unpaired *t*-test. \**P* < 0.05, \*\**P* < 0.01, \*\*\**P* < 0.001, ns = not significant.

**Supplementary Figure 2:**
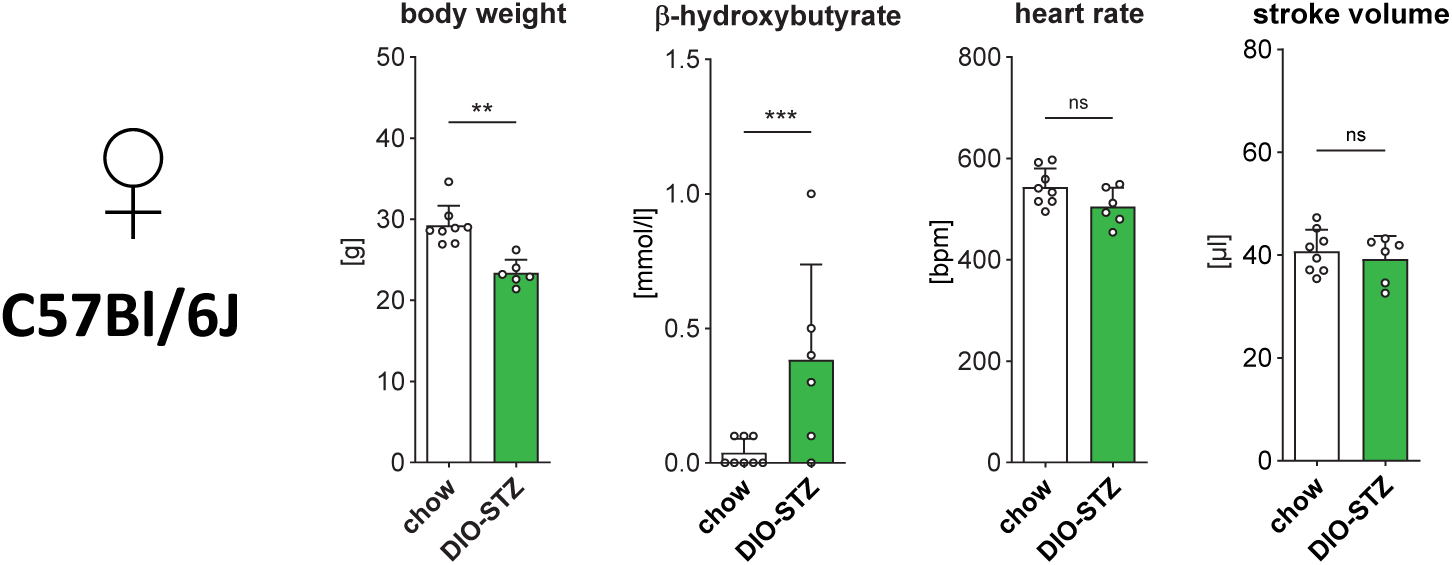
Additional data from phenotyping of DIO-STZ treatment in female C57Bl/6J mice. Summarized data of body weight, β-hydroxybutyrate plasma levels, heart rate during echocardiographic analysis, and stroke volume determined by echocardiography are shown (n=6-8 per group). Data are shown as means ± SD. Statistical analysis was performed using unpaired t-test. \*\**P* < 0.01, \*\*\**P* < 0.001, ns = not significant.

**Supplementary Figure 3:**
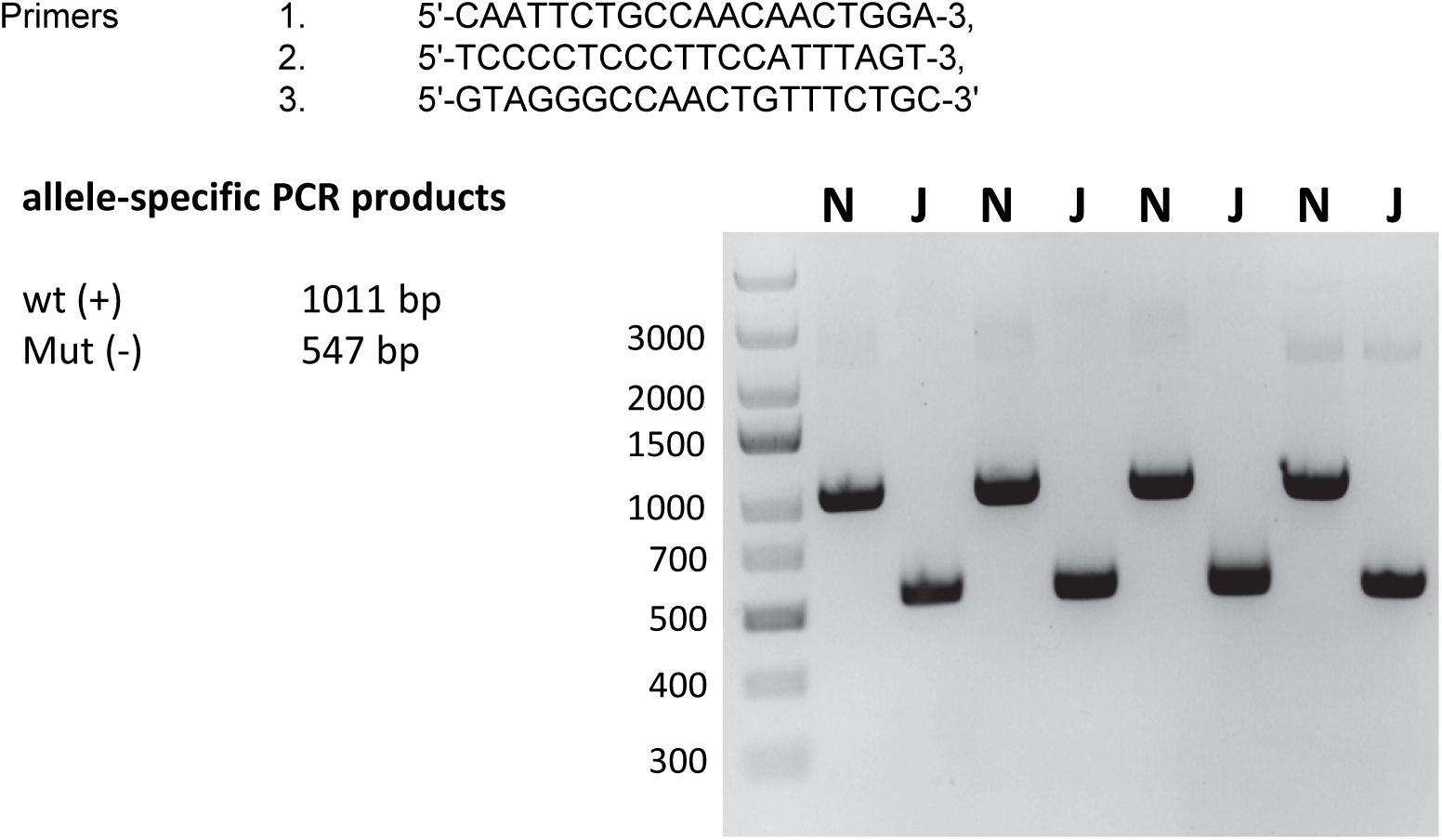
Genotyping of *Nnt* mutation. The presence of the *Nnt* mutation in C57Bl/6J mice (Mut; J) was confirmed by PCR analysis using indicated primers. C57Bl/6N mice were used as positive controls (wt; N).

**Supplementary Figure 4:**
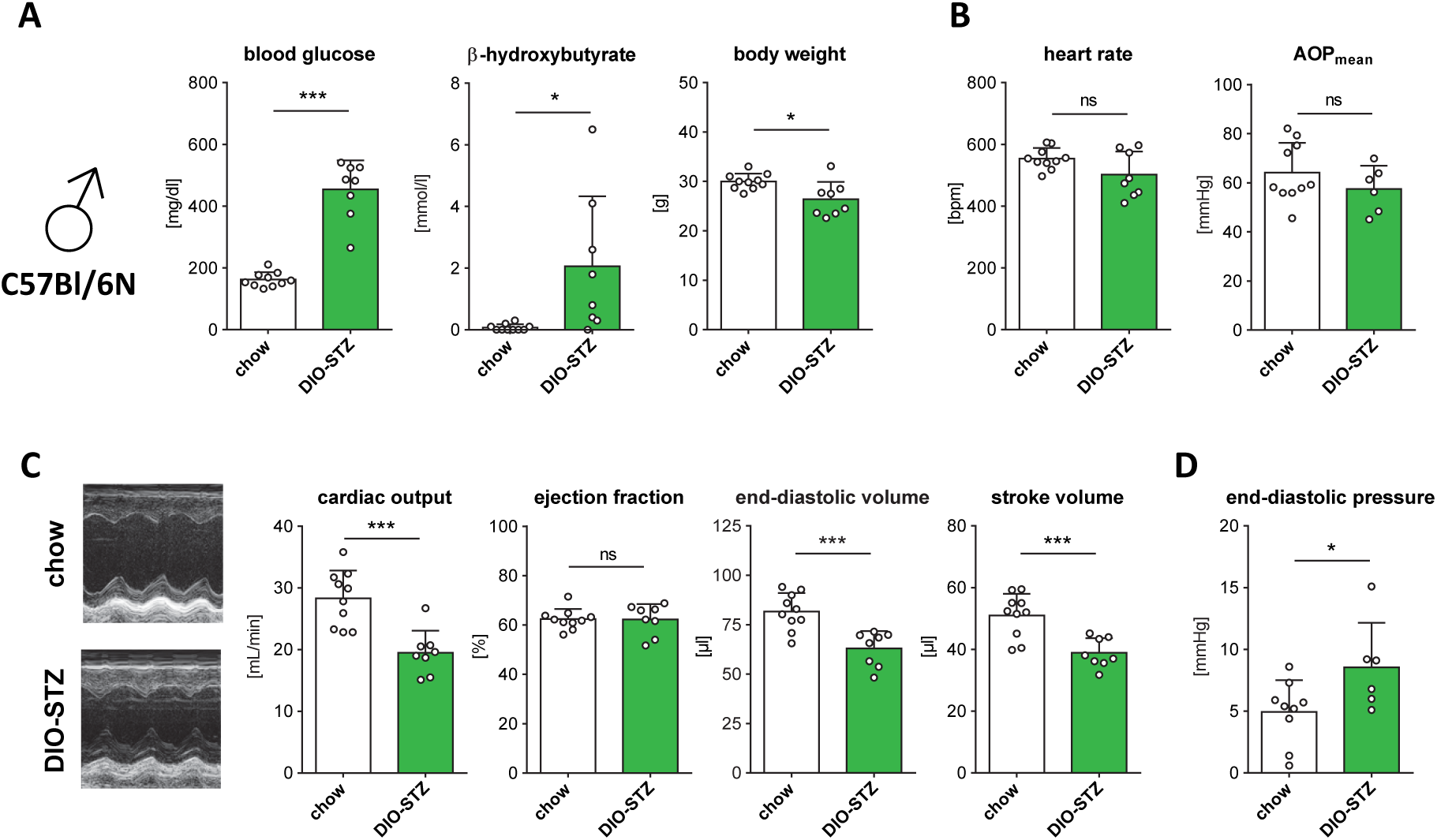
Phenotyping of DIO-STZ treatment in male C57Bl/6N mice. (A) Summarized data of blood glucose, β-hydroxybutyrate plasma levels, and body weight (n=8-9). (B) Summarized data of heart rate during echocardiographic analysis (n=8-10), and mean aortic pressure (AOP_mean_; n=6-10). (C) Representative echocardiographic SAX m-mode images of male C57Bl/6N chow and DIO-STZ hearts (left), and summarized data for cardiac output, ejection fraction, end-diastolic volume, and stroke volume (n=8- 10). (D) Summarized data of end-diastolic pressure obtained by left ventricular catheterization (n=6-9). Data are shown as means ± SD. Statistical analysis was performed using unpaired t-test. \**P* < 0.05, \*\**P* < 0.01, \*\*\**P* < 0.001, ns = not significant.

**Supplementary Table S1:**
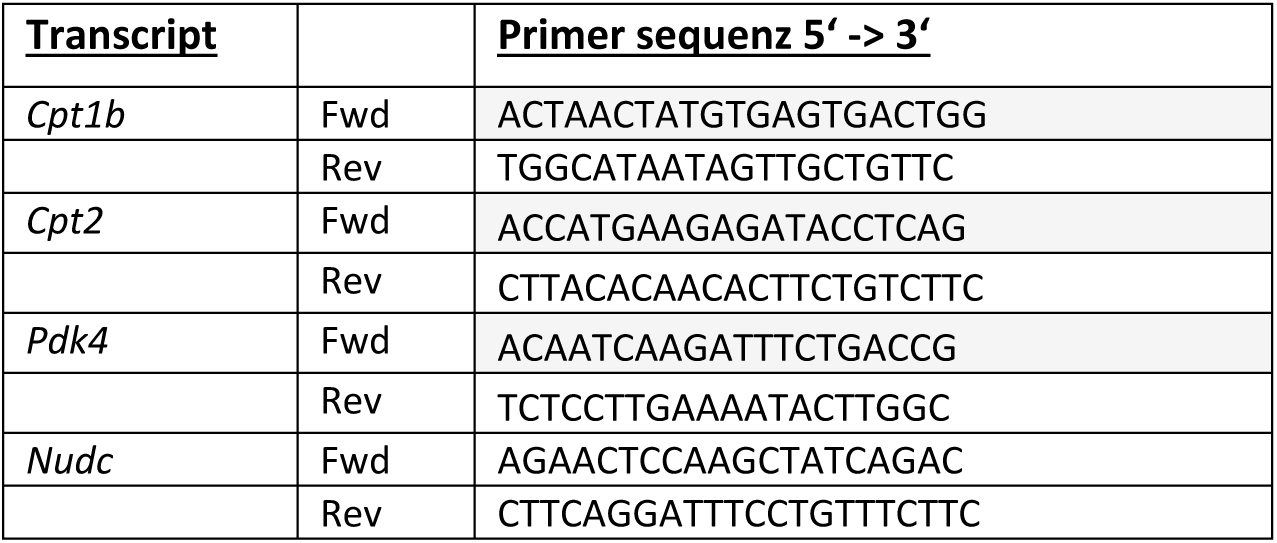
qPCR primers.

**Supplementary Table S2:** Blood cell count of chow and DIO-STZ mice.

| Parameter | Unit | chow | DIO-STZ | p-value |
| --- | --- | --- | --- | --- |
| *Red blood cell count* |  |  |  |  |
| Erythrocytes | $10^{12}/l$ | $9.33 \pm 0.37$ | $9.85 \pm 0.29$ | 0.023 |
| HGB | g/dl | $12.8 \pm 0.7$ | $13.7 \pm 0.4$ | 0.016 |
| HCT | % | $39.3 \pm 1.2$ | $41.8 \pm 1.1$ | 0.004 |
| *Red blood cell indices* |  |  |  |  |
| MCV | fl | $42.2 \pm 1.2$ | $42.2 \pm 0.4$ | 1.000 |
| MCH | pg | $13.7 \pm 0.7$ | $14.0 \pm 0.3$ | 0.420 |
| MCHC | g/dl | $32.5 \pm 1.1$ | $32.9 \pm 1.9$ | 0.575 |
| *White blood cell count* |  |  |  |  |
| Leukocytes | $10^9/l$ | $3.59 \pm 1.8$ | $3.58 \pm 1.25$ | 0.991 |
| Lymphocytes | $10^9/l$ | $2.67 \pm 0.79$ | $3.29 \pm 1.19$ | 0.310 |
| Monocytes | $10^9/l$ | $0.17 \pm 0.18$ | $0.09 \pm 0.05$ | 0.288 |
| Neutrophils | $10^9/l$ | $0.75 \pm 0.97$ | $0.20 \pm 0.08$ | 0.195 |
| *Platelet count* |  |  |  |  |
| Platelets | $10^9/l$ | $526 \pm 60$ | $607 \pm 360$ | 0.597 |

HGB, hemoglobin; HCT, hematocrit; MCV, mean corpuscular volume; MCH, mean corpuscular hemoglobin; MCHC, mean corpuscular hemoglobin concentration; Data are mean ± SD; unpaired t-test. n=6 per group.

## References

1. International Diabetes Federation. IDF Diabetes Atlas. 10th ed. Brussels, Belgium2021.

2. Holman RR, Paul SK, Bethel MA, Matthews DR, Neil HA. 10-year follow-up of intensive glucose control in type 2 diabetes. N Engl J Med. 2008;359(15):1577–89.

3. Dunlay SM, Givertz MM, Aguilar D, Allen LA, Chan M, Desai AS, et al. Type 2 Diabetes Mellitus and Heart Failure, A Scientific Statement From the American Heart Association and Heart Failure Society of America. J Card Fail. 2019;25(8):584–619.

4. McAllister DA, Read SH, Kerssens J, Livingstone S, McGurnaghan S, Jhund P, et al. Incidence of Hospitalization for Heart Failure and Case-Fatality Among 3.25 Million People With and Without Diabetes Mellitus. Circulation. 2018;138(24):2774–86.

5. Nichols GA, Gullion CM, Koro CE, Ephross SA, Brown JB. The incidence of congestive heart failure in type 2 diabetes: an update. Diabetes Care. 2004;27(8):1879–84.

6. Park JJ. Epidemiology, Pathophysiology, Diagnosis and Treatment of Heart Failure in Diabetes. Diabetes Metab J. 2021;45(2):146–57.

7. Teramoto K, Teng TK, Chandramouli C, Tromp J, Sakata Y, Lam CS. Epidemiology and Clinical Features of Heart Failure with Preserved Ejection Fraction. Card Fail Rev. 2022;8:e27.

8. Vasan RS, Xanthakis V, Lyass A, Andersson C, Tsao C, Cheng S, et al. Epidemiology of Left Ventricular Systolic Dysfunction and Heart Failure in the Framingham Study: An Echocardiographic Study Over 3 Decades. JACC Cardiovasc Imaging. 2018;11(1):1–11.

9. Dalos D, Mascherbauer J, Zotter-Tufaro C, Duca F, Kammerlander AA, Aschauer S, et al. Functional Status, Pulmonary Artery Pressure, and Clinical Outcomes in Heart Failure With Preserved Ejection Fraction. J Am Coll Cardiol. 2016;68(2):189–99.

10. Obokata M, Reddy YNV, Pislaru SV, Melenovsky V, Borlaug BA. Evidence Supporting the Existence of a Distinct Obese Phenotype of Heart Failure With Preserved Ejection Fraction. Circulation. 2017;136(1):6–19.

11. Anker SD, Butler J, Filippatos G, Ferreira JP, Bocchi E, Bohm M, et al. Empagliflozin in Heart Failure with a Preserved Ejection Fraction. N Engl J Med. 2021;385(16):1451–61.

12. Kosiborod MN, Abildstrom SZ, Borlaug BA, Butler J, Rasmussen S, Davies M, et al. Semaglutide in Patients with Heart Failure with Preserved Ejection Fraction and Obesity. N Engl J Med. 2023;389(12):1069–84.

13. Reed MJ, Meszaros K, Entes LJ, Claypool MD, Pinkett JG, Gadbois TM, et al. A new rat model of type 2 diabetes: the fat-fed, streptozotocin-treated rat. Metabolism. 2000;49(11):1390–4.

14. Skovso S. Modeling type 2 diabetes in rats using high fat diet and streptozotocin. J Diabetes Investig. 2014;5(4):349–58.

15. Heinen A, Nederlof R, Panjwani P, Spychala A, Tschaidse T, Reffelt H, et al. IGF1 Treatment Improves Cardiac Remodeling after Infarction by Targeting Myeloid Cells. Mol Ther. 2019;27(1):46–58.

16. Heinen A, Raupach A, Behmenburg F, Holscher N, Flogel U, Kelm M, et al. Echocardiographic Analysis of Cardiac Function after Infarction in Mice: Validation of Single-Plane Long-Axis View Measurements and the Bi-Plane Simpson Method. Ultrasound Med Biol. 2018;44(7):1544–55.

17. Pieske B, Tschope C, de Boer RA, Fraser AG, Anker SD, Donal E, et al. How to diagnose heart failure with preserved ejection fraction: the HFA-PEFF diagnostic algorithm: a consensus recommendation from the Heart Failure Association (HFA) of the European Society of Cardiology (ESC). Eur Heart J. 2019;40(40):3297–317.

18. Schnelle M, Catibog N, Zhang M, Nabeebaccus AA, Anderson G, Richards DA, et al. Echocardiographic evaluation of diastolic function in mouse models of heart disease. J Mol Cell Cardiol. 2018;114:20–8.

19. Thomsen R, Solvsten CA, Linnet TE, Blechingberg J, Nielsen AL. Analysis of qPCR data by converting exponentially related Ct values into linearly related X0 values. J Bioinform Comput Biol. 2010;8(5):885–900.

20. Bottermann K, Spychala A, Eliacik A, Amin E, Moussavi-Torshizi SE, Klocker N, et al. Extracellular flux analysis in intact cardiac tissue slices-A novel tool to investigate cardiac substrate metabolism in mouse myocardium. Acta Physiol (Oxf). 2023;239(2):e14004.

21. Heinen A, Aldakkak M, Stowe DF, Rhodes SS, Riess ML, Varadarajan SG, et al. Reverse electron flow- induced ROS production is attenuated by activation of mitochondrial Ca^2+^-sensitive K^+^ channels. Am J Physiol Heart Circ Physiol. 2007;293(3):H1400–7.

22. Bradford MM. A rapid and sensitive method for the quantitation of microgram quantities of protein utilizing the principle of protein-dye binding. Anal Biochem. 1976;72:248–54.

23. Pepin ME, Konrad PJM, Nazir S, Bazgir F, Maack C, Nickel A, et al. Mitochondrial NNT Promotes Diastolic Dysfunction in Cardiometabolic HFpEF. Circ Res. 2025;136(12):1564–78.

24. Borlaug BA, Jensen MD, Kitzman DW, Lam CSP, Obokata M, Rider OJ. Obesity and heart failure with preserved ejection fraction: new insights and pathophysiological targets. Cardiovasc Res. 2023;118(18):3434–50.

25. Kitzman DW, Lam CSP. Obese Heart Failure With Preserved Ejection Fraction Phenotype: From Pariah to Central Player. Circulation. 2017;136(1):20–3.

26. Conceicao G, Heinonen I, Lourenco AP, Duncker DJ, Falcao-Pires I. Animal models of heart failure with preserved ejection fraction. Neth Heart J. 2016;24(4):275–86.

27. Valero-Munoz M, Backman W, Sam F. Murine Models of Heart Failure with Preserved Ejection Fraction: a “Fishing Expedition”. JACC Basic Transl Sci. 2017;2(6):770–89.

28. Withaar C, Lam CSP, Schiattarella GG, de Boer RA, Meems LMG. Heart failure with preserved ejection fraction in humans and mice: embracing clinical complexity in mouse models. Eur Heart J. 2021;42(43):4420–30.

29. Rosas PC, Neves LAA, Patel N, Tran D, Pereira CH, Bonilla KR, et al. Early pathological mechanisms in a mouse model of heart failure with preserved ejection fraction. Am J Physiol Heart Circ Physiol. 2024;327(6):H1524–H43.

30. Daou D, Tong D, Schiattarella GG, Gillette TG, Hill JA. What is Cardiometabolic HFpEF and How Can We Study it Preclinically? JACC Basic Transl Sci. 2025:101295.

31. Velagic A, Li M, Deo M, Li JC, Kiriazis H, Donner DG, et al. A high-sucrose diet exacerbates the left ventricular phenotype in a high fat-fed streptozotocin rat model of diabetic cardiomyopathy. Am J Physiol Heart Circ Physiol. 2023;324(2):H241–H57.

32. Guan Y, Zhang M, Lacy C, Shah S, Epstein FH, Yan Z. Endurance Exercise Training Mitigates Diastolic Dysfunction in Diabetic Mice Independent of Phosphorylation of Ulk1 at S555. Int J Mol Sci. 2024;25(1).

33. Wu XQ, Liu H, Brooks A, Xu SW, Luo JQ, Steiner R, et al. SIRT6 Mitigates Heart Failure With Preserved Ejection Fraction in Diabetes. Circulation Research. 2022;131(11):926–43.

34. Sun Q, Guven B, Wagg CS, Almeida de Oliveira A, Silver H, Zhang L, et al. Mitochondrial fatty acid oxidation is the major source of cardiac adenosine triphosphate production in heart failure with preserved ejection fraction. Cardiovasc Res. 2024;120(4):360–71.

35. Sun Q, Wagg CS, Wong N, Wei K, Ketema EB, Zhang L, et al. Alterations of myocardial ketone metabolism in heart failure with preserved ejection fraction (HFpEF). ESC Heart Fail. 2025.

36. Murashige D, Jang C, Neinast M, Edwards JJ, Cowan A, Hyman MC, et al. Comprehensive quantification of fuel use by the failing and nonfailing human heart. Science. 2020;370(6514):364–8.

37. Schiattarella GG, Altamirano F, Tong D, French KM, Villalobos E, Kim SY, et al. Nitrosative stress drives heart failure with preserved ejection fraction. Nature. 2019;568(7752):351–6.

38. Pandey A, Omar W, Ayers C, LaMonte M, Klein L, Allen NB, et al. Sex and Race Differences in Lifetime Risk of Heart Failure With Preserved Ejection Fraction and Heart Failure With Reduced Ejection Fraction. Circulation. 2018;137(17):1814–23.

39. Beale AL, Meyer P, Marwick TH, Lam CSP, Kaye DM. Sex Differences in Cardiovascular Pathophysiology: Why Women Are Overrepresented in Heart Failure With Preserved Ejection Fraction. Circulation. 2018;138(2):198–205.

40. Duca F, Zotter-Tufaro C, Kammerlander AA, Aschauer S, Binder C, Mascherbauer J, et al. Gender-related differences in heart failure with preserved ejection fraction. Sci Rep. 2018;8(1):1080.

41. Cao Y, Vergnes L, Wang YC, Pan C, Chella Krishnan K, Moore TM, et al. Sex differences in heart mitochondria regulate diastolic dysfunction. Nat Commun. 2022;13(1):3850.

42. Tong D, Schiattarella GG, Jiang N, May HI, Lavandero S, Gillette TG, et al. Female Sex Is Protective in a Preclinical Model of Heart Failure With Preserved Ejection Fraction. Circulation. 2019;140(21):1769–71.

